# A Framework for Intelligence and Cortical Function Based on Grid Cells in the Neocortex

**DOI:** 10.1101/442418

**Authors:** Jeff Hawkins, Marcus Lewis, Mirko Klukas, Scott Purdy, Subutai Ahmad

## Abstract

How the neocortex works is a mystery. In this paper we propose a novel framework for understanding its function. Grid cells are neurons in the entorhinal cortex that represent the location of an animal in its environment. Recent evidence suggests that grid cell-like neurons may also be present in the neocortex. We propose that grid cells exist throughout the neocortex, in every region and in every cortical column. They define a location-based framework for how the neocortex functions. Whereas grid cells in the entorhinal cortex represent the location of one thing, the body relative to its environment, we propose that cortical grid cells simultaneously represent the location of many things. Cortical columns in somatosensory cortex track the location of tactile features relative to the object being touched and cortical columns in visual cortex track the location of visual features relative to the object being viewed. We propose that mechanisms in the entorhinal cortex and hippocampus that evolved for learning the structure of environments are now used by the neocortex to learn the structure of objects. Having a representation of location in each cortical column suggests mechanisms for how the neocortex represents object compositionality and object behaviors. It leads to the hypothesis that every part of the neocortex learns complete models of objects and that there are many models of each object distributed throughout the neocortex. The similarity of circuitry observed in all cortical regions is strong evidence that even high-level cognitive tasks are learned and represented in a location-based framework.

## Introduction

The human neocortex learns an incredibly complex and detailed model of the world. Each of us can recognize thousands of objects. We know how these objects appear through vision, touch, and audition, we know how these objects behave and change when we interact with them, and we know their location in the world. The human neocortex also learns models of abstract objects, structures that don’t physically exist or that we cannot directly sense. The circuitry of the neocortex is also complex. Understanding how the complex circuitry of the neocortex learns complex models of the world is one of the primary goals of neuroscience.

Vernon Mountcastle was the first to propose that all regions of the neocortex are fundamentally the same. What distinguishes one region from another, he argued, is mostly determined by the inputs to a region and not by differences in intrinsic circuitry and function. He further proposed that a small volume of cortex, a cortical column, is the unit of replication (Mountcastle, 1978). These are compelling ideas, but it has been difficult to identify what a column could do that is sufficient to explain all cognitive abilities. Today, the most common view is that the neocortex processes sensory input in a series of hierarchical steps, extracting more and more complex features until objects are recognized. Although this view explains some aspects of sensory inference, it fails to explain the richness of human behavior, how we learn multidimensional models of objects, and how we learn how objects themselves change and behave when we interact with them. It also fails to explain what most of the circuitry of the neocortex is doing. In this paper we propose a new theoretical framework based on location processing that addresses many of these shortcomings.

Over the past few decades some of the most exciting advances in neuroscience have been related to “grid cells” and “place cells”. These neurons exist in the hippocampal complex of mammals, a set of regions, which, in humans, is roughly the size and shape of a finger, one on each side of the brain. Grid cells in combination with place cells learn maps of the world (Hafting et al., 2005; Moser et al., 2008; O’Keefe and Dostrovsky, 1971). Grid cells represent the current location of an animal relative to those maps. Modeling work on the hippocampus has demonstrated the power of these neural representations for episodic and spatial memory (Byrne et al., 2007; Hasselmo, 2012; Hasselmo et al., 2010), and navigation (Erdem and Hasselmo, 2014; Bush et al., 2015). There is also evidence that grid cells play a role in more abstract cognitive tasks (Behrens et al., 2018; Constantinescu et al., 2016).

Recent experimental evidence suggests that grid cells may also be present in the neocortex. Using fMRI, (Doeller et al., 2010; Constantinescu et al., 2016; Julian et al., 2018) have found signatures of grid cell-like firing patterns in prefrontal and parietal areas of the neocortex. Using single cell recording in humans, (Jacobs et al., 2013) have found more direct evidence of grid cells in frontal cortex. (Long and Zhang, 2018), using multiple tetrode recordings, have reported finding cells exhibiting grid cell, place cell, and conjunctive cell responses in rat S1. Our team has proposed that prediction of sensory input by the neocortex requires a representation of an object-centric location to be present throughout the sensory regions of the neocortex, which is consistent with grid cell-like mechanisms (Hawkins et al., 2017).

Here we propose that grid cell-like neurons exist in every column of the neocortex. Whereas grid cells in the medial entorhinal cortex (MEC) primarily represent the location of one thing, the body, we suggest that cortical grid cells simultaneously represent the location of multiple things. Columns in somatosensory cortex that receive input from different parts of the body represent the location of those inputs in the external reference frames of the objects being touched. Similarly, cortical columns in visual cortex that receive input from different patches of the retinas represent the location of visual input in the external reference frames of the objects being viewed. Whereas grid cells and place cells learn models of environments via movement of the body, we propose that cortical grid cells combined with sensory input learn models of objects via movement of the sensors.

Although much is known about the receptive field properties of grid cells in MEC and how these cells encode location (Rowland et al., 2016), the underlying mechanisms leading to those properties is not known. Experimental results suggest that grid cells have unique membrane and dendritic properties (Domnisoru et al., 2013; Schmidt-Hieber et al., 2017). There are two leading computational candidates, oscillatory interference models (O’Keefe and Burgess, 2005; Burgess et al., 2007; Giocomo et al., 2007; Burgess, 2008; Giocomo et al., 2011) and continuous attractor models (Fuhs and Touretzky, 2006; Burak and Fiete, 2009). The framework proposed in this paper assumes that “cortical grid cells” exhibit similar physiological properties as grid cells in MEC, but the framework is not dependent on how those properties arise.

Throughout this paper we refer to “cortical columns”. We use this term similarly to Mountcastle, to represent a small area of neocortex that spans all layers in depth and of sufficient lateral extent to capture all cell types and receptive field responses. For this paper, a cortical column is not a physically demarked entity. It is a convenience of nomenclature. We typically think of a column as being about one square millimeter of cortical area, although this size is not critical and could vary by species and region.

## How Grid Cells Represent Location

To understand our proposal, we first review how grid cells in the entorhinal cortex are believed to represent space and location, **Figure 1**. Although many details of grid cell function remain unknown, general consensus exists on the following principles. A grid cell is a neuron that becomes active at multiple locations in an environment, typically in a grid-like, or tiled, triangular lattice. A “grid cell module” is a set of grid cells that activate with the same lattice spacing and orientation but at shifted locations within an environment. As an animal moves, the active grid cells in a grid cell module change to reflect the animal’s updated location. This change occurs even if the animal is in the dark, telling us that grid cells are updated using an internal, or “efference”, copy of motor commands (Kropff et al., 2015; McNaughton et al., 2006; Moser et al., 2008). This process, called “path integration”, has the desirable property that regardless of the path of movement, when the animal returns to the same physical location, then the same grid cells in a module will be active.

**Figure 1.**
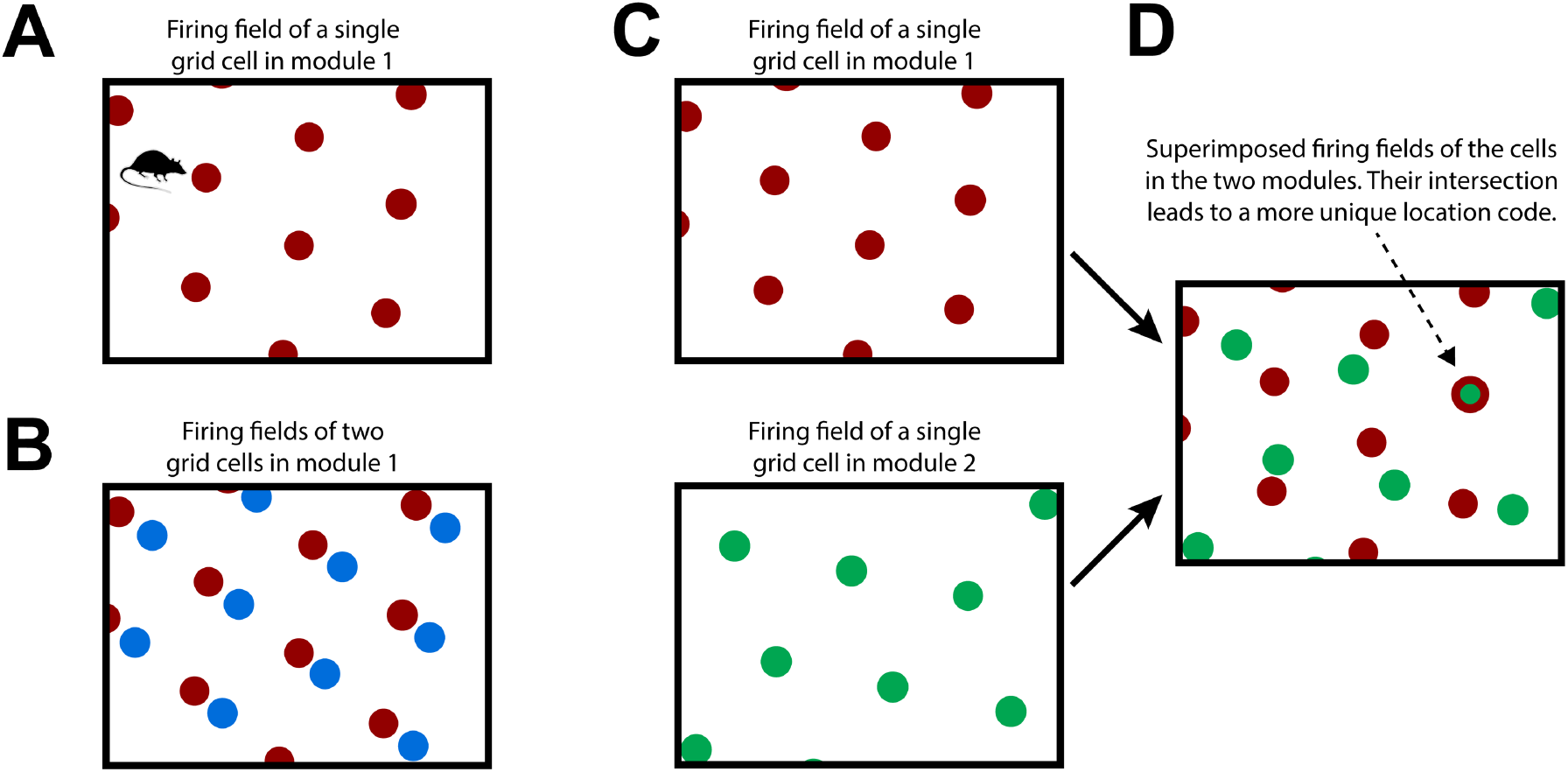
How Grid Cells Represent Location. **(A)** An individual grid cell becomes active at multiple locations (red circles) as an animal moves about an environment (rectangle). The locations of activation form a periodic grid-like lattice. The activation locations are always the same for any particular environment. **(B)** A grid cell module is a set of grid cells that activate at the same spacing and orientation but at different positions in the environment. The activation locations for two grid cells in a grid cell module are shown (red and blue dots). Every location in an environment will activate one or more grid cells in a module. Because of the periodic activation of grid cells, a single grid cell module cannot represent unique locations. **(C)** Multiple grid cell modules (two shown, top and bottom) tile the same space at different orientations and/or spacings. **(D)** Although a single module cannot represent unique locations in an environment, the activity across multiple modules can. This rectangle shows the superimposed firing fields of the two grid cells from C). Note that when the two cells (red and green) fire together, only one location is possible (indicated by arrow). The number of locations that can be represented increases exponentially with the number of modules.

Due to tiling, a single grid cell module cannot represent a unique location. To form a representation of a unique location requires looking at the active cells in multiple grid cell modules where each grid cell module differs in its tile spacing and/or orientation relative to the environment, **Figure 1C and 1D**. For example, if a single grid cell module can represent twenty different locations before repeating, then ten grid cell modules can represent approximately 20^10^ different locations before repeating. (Fiete et al., 2008) This method of representing location has several desirable properties.

### 1) Large representational capacity

The number of locations that can be represented by a set of grid cell modules is large as it scales exponentially with the number of modules.

### 2) Path integration works from any location

No matter what location the network starts with, path integration will work. This is a form of generalization. The path integration properties have to be learned once for each grid cell module, but then apply to all locations, even those the animal has never been in before.

### 3) Locations are unique to each environment

Every learned environment is associated with a set of unique locations. Experimental recordings suggest that upon entering a learned environment, entorhinal grid cell modules “anchor” differently (Marozzi et al., 2015; Rowland and Moser, 2014). (The term “anchor” refers to selecting which grid cells in each module should be active at the current location.) This suggests that the current location and all the locations that the animal can move to in that environment will, with high certainty, have representations that are unique to that environment (Fiete et al., 2008; Sreenivasan and Fiete, 2011).

Combining these properties, we can now broadly describe how grid cells represent an environment such as a room, **Figure 2A**. An environment consists of a set of location representations that are related to each other via path integration (i.e. the animal can move between these location representations). Each location representation in the set is unique to that environment and will not appear in any other environment. An environment consists of all the locations that the animal can move among, including locations that have not been visited, but could be visited. Associated with some of the location representations are observable landmarks.

**Figure 2.**
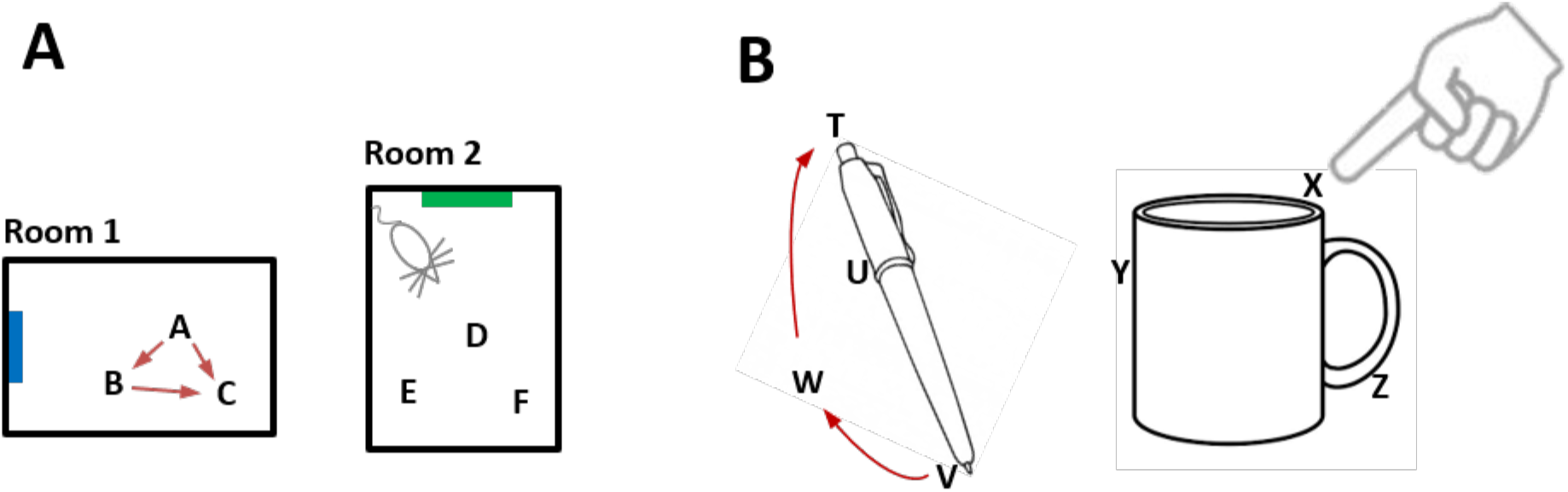
Representing Objects as Location Spaces. We propose that the neocortex learns the structure of objects in the same way that the entorhinal cortex and hippocampus learn the structure of environments. **(A)** Two rooms that a rodent has learned. Because of distinct landmarks (suggested by blue and green rectangles) an animal will perceive these as different rooms. Locations in a room are represented by the activity in a set of grid cell modules in the entorhinal cortex. Three locations are shown for each room (A,B,C and D,E,F). Representations of location are unique to both the location in a room and the room. Therefore, if an animal can determine it is in location A, then it knows what room it is in (Room1) and its location in the room. The locations associated with a room are united via movement and path integration. As an animal moves, the representation of location is updated (red arrows) based on an internal copy of its motor behavior. By exploring a room, the animal learns the features associated with locations in the room. **(B)** We propose that objects such as a pen or coffee cup are similarly defined by a set of locations (four labeled for the pen and three labeled for the cup). Grid cells in the neocortex represent the location of a sensor patch (for example, tip of finger) in the location space of the object. Locations in an object’s space are unique to the object and the location relative to the object. An object’s space includes locations that can be moved to but don’t necessarily have an associated feature. For example, location W is part of the pen because a finger can move from V to W to T via path integration. By moving and exploring the object, the neocortex learns the features associated with locations of the object.

## Grid Cells In The Neocortex

Now let us consider a patch of neocortex that receives input from the tip of a finger, **Figure 2B**. Our proposal is that some of the neurons in that patch of cortex represent the location of the fingertip as it explores an object. When the finger moves, these cortical grid cells update their representation of location via a motor efference copy and path integration. Objects, such as a coffee cup, have an associated set of locations, in the same way that environments, such as a room, have an associated set of locations. Associated with some of the object’s locations are observable features. The cortical area receiving input from the fingertip tracks the location of the sensory input from the fingertip in the location space of the object. Through movement and sensation, the fingertip cortical area learns models of objects in the same way that grid cells and place cells learn models of environments. Whereas the entorhinal cortex tracks the location of the body, different areas of the neocortex independently track the location of each movable sensory patch. For example, each area of somatosensory cortex tracks the location of sensory input from its associated body part. These areas operate in parallel and build parallel models of objects. The same basic method applies to vision. Patches of the retina are analogous to patches of skin. Different parts of the retina observe different locations on an object. Each patch of cortex receiving visual input tracks the location of its visual input in the location space of the object being observed. As the eyes move, visual cortical columns sense different locations on an object and learn parallel models of the observed object.

We have now covered the most basic aspects of our proposal.

1) Every cortical column has neurons that perform a function similar to grid cells. The activation pattern of these cortical grid cells represents the location of the column’s input relative to an external reference frame. The location representation is updated via a motor efference copy and path integration.

2) Cortical columns learn models of objects in the world similarly to how grid cells and place cells learn models of environments. The models learned by cortical columns consist of a set of location representations that are unique to each object, and where some of the locations have observable features.

## A Location-Based Framework For Cortical Computation

Our proposal suggests that cortical columns are more powerful than previously assumed. By pairing input with a grid cell-derived representation of location, individual columns can learn complex models of structure in the world (see also (Lewis et al., 2018)). In this section we show how a location-based framework allows neurons to learn the rich models that we know the neocortex is capable of.

### Object Compositionality

Objects are composed of other objects arranged in a particular way. For example, it would be inefficient to learn the morphology of a coffee cup by remembering the sensory sensation at each location on the cup. It is far more efficient to learn the cup as the composition of previously learned objects, such as a cylinder and a handle. Consider a coffee cup with a logo on it, **Figure 3A**. The logo exists in multiple places in the world and is itself a learned “object”. To represent the cup with the logo we need a way of associating one object, “the logo”, at a relative position to another object, “the cup”. Compositional structure is present in almost all objects in the world, therefore cortical columns must have a neural mechanism that represents a new object as an arrangement of previously-learned objects. How can this functionality be achieved?

**Figure 3.**
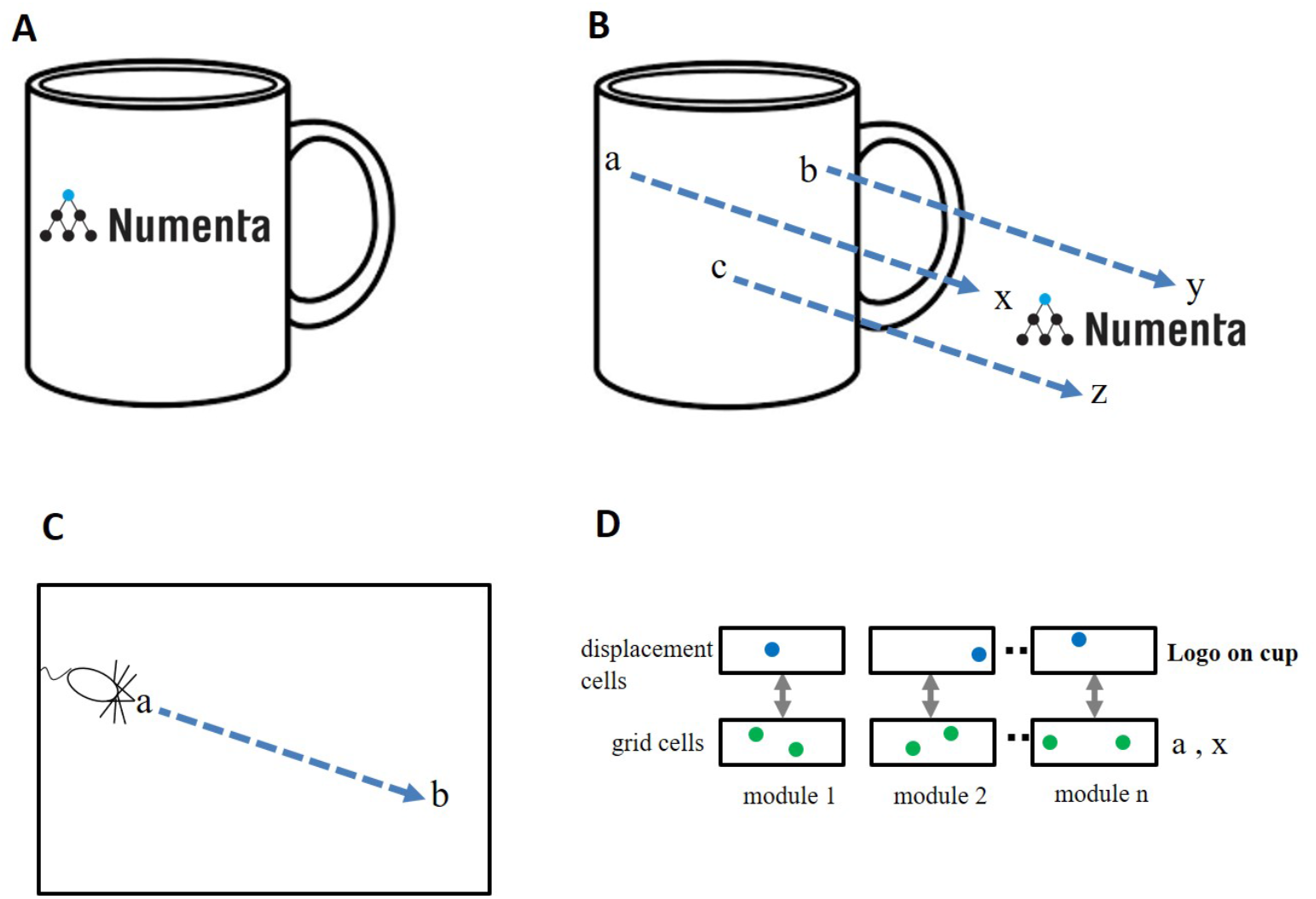
Representing Objects as Compositions of other Objects. **(A)** The neocortex can learn an object, such as a “coffee cup with logo”, as a composition of two previously learned objects, “cup” and “logo”. The goal is to represent this relationship efficiently, without any relearning. **(B)** The cup and the logo each have their own unique set of locations. Three locations are shown in cup space (**a, b, c**) and three locations are shown in logo space (**x, y, z**). When the logo is placed on the cup there is a fixed one-to-one mapping between locations in cup space and locations in logo space. This relationship can be represented as a displacement vector between the two spaces (blue arrows). **(C)** Animals exploring an environment can determine the direction and distance from their current location, **a**, to a previously visited target location, **b**, even if they have never taken this path before. Determining the displacement between two locations in the same space (e.g. **a** to **b** in panel C) is equivalent to determining the displacement between two locations in separate spaces (e.g. **a** to **x** in panel B). **(D)** A method to determine the displacement between two locations. Each grid cell module is paired with a displacement cell module. Cells in a displacement cell module (blue dots) respond to a particular displacement between pairs of grid cells (green dots). Any two pairs of grid cells with the same displacement in physical space will activate the same displacement cell. Displacement cells cannot represent a unique displacement in the same way that grid cells cannot represent a unique location. However, the set of active cells in multiple displacement cell modules (three shown) will represent a unique displacement. Because the set of active grid cells in multiple grid cell modules is unique to objects (cup and logo), the set of active displacement cells will also be unique (to both the cup and logo). Thus, a set of active displacement cells can represent the relative placement of two specific objects (location of logo on cup).

We have proposed that each object is associated with a set of locations which are unique to the object and comprise a space around the object. If a finger is touching the coffee cup with the logo, then the cortical grid cells representing the location of the finger can at one moment represent the location of the finger in the space of the coffee cup and at another moment, after re-anchoring, represent the location of the finger in the space of the logo. If the logo is attached to the cup, then there is a fixed, one-to-one, relationship between any point in the space of the logo and the equivalent point in the space of the cup, **Figure 3B**. The task of representing the logo on the cup can be achieved by creating a “displacement” vector that converts any point in cup space to the equivalent point in logo space.

Determining the displacement between two objects is similar to a previously-studied navigation problem, specifically, how an animal knows how to get from point **a** to point **b** within an environment, **Figure 3C**. Mechanisms that solve the navigation problem (determining the displacement between two points in the same space) can also solve the object composition problem (determining the displacement between two points in two different spaces).

### Displacement Cells

Several solutions have been proposed for solving the point-to-point navigation problem using grid cells. One class of solutions detects the difference between two sets of active grid cells across multiple grid cell modules (Bush et al., 2015) and another uses linear look-ahead probes using grid cells for planning and computing trajectories (Erdem and Hasselmo, 2014). We suggest an alternate but related solution. Our proposal also relies on detecting differences between two sets of active grid cells, however, we propose this is done on a grid cell module by grid cell module basis. We refer to these cells as “displacement cells”. (See **Box 1** for a more thorough description.) Displacement cells are similar to grid cells in that they can’t on their own represent a unique displacement. (In the example of Box 1, a displacement cell that represents a displacement of “two to the right and one up”, would also be active for “five over and four up”.) However, the cell activity in multiple displacement cell modules represents a unique displacement in much the same way as the cell activity in multiple grid cell modules represents a unique location, **Figure 3D**. Hence, a single displacement vector can represent the logo on the coffee cup at a specific relative position. Note, a displacement vector not only represents the relative position of two objects, it also is unique to the two objects. Complex objects can be represented by a set of displacement vectors which define the components of an object and how they are arranged relative to each other. This is a highly efficient means of representing and storing the structure of objects.

#### Box 1: Displacement Cells

Here we describe a possible network mechanism for a layer of displacement cell modules, derived from a layer of grid cell modules. We first illustrate the mechanism for a single module, and then describe multiple modules.

In the lower box of **Figure 1A**, we show the cells in a single grid cell module. In this simple example there are 9 cells, arranged in a 3X3 lattice. For convenience we assign (x,y) coordinates to each cell where x and y can be 0, 1, or 2. The cells in this lattice are arranged such that neighboring cells represent neighboring positions. For example, if cell *G*_1,1_ is active for a location on an object, a movement one step to the right will cause step to the right will cause *G*_2,1_ to become active. Since the cells tile space, an additional movement one step to the right will cause *G*_0,1_ to become active.

Each grid cell module is paired with a displacement module (upper box of **Figure 1A**). Each cell in the displacement module corresponds to a particular relative movement. It becomes active if any of the grid cell pairs corresponding to its displacement become active in the appropriate order. For example, cell *D*_1,0_, representing a displacement one step to the right, would become active if grid cell *G*_2,2_ becomes active after grid cell *G*_1,2_. It would also become active if cell *G*_1,1_ becomes active after *G*_0,1_, etc. A possible neural implementation is that these pairs of grid cells connect to independent dendritic segments on cell *D*_1,0_ such that any of these transitions cause *D*_1,0_ to become active.

A grid cell module can be thought of as implementing a residue number system (Omondi and Premkumar, 2007; Sreenivasan and Fiete, 2011). Viewed in this light a displacement module implements residual subtraction operations. If the active cell moves from location *G*_*x*1,*y1*_ to *G*_*x*2,*y*2_, the displacement cell in position ((*x*2 – *x*1) *modulo* 3, (*y*2 – *y*1) *modulo* 3) will become active. Each cell in the displacement module connects to 9 pairs of grid cells. Cell *D*_1,0_ thus becomes active for any of the following 9 pairs of cells:

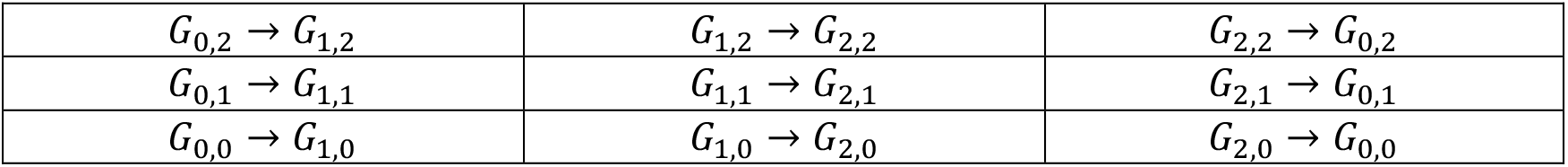

**Figure 1B** shows all the transitions within the grid cell module that cause *D*_1,0_ to become active. These transitions form the input to the displacement cells.

The number of possible displacements is tied to the number of cells in a grid cell module. Note that the magnitude and direction of a shift is ambiguous within a module. You cannot distinguish a displacement two steps to the right from a displacement one step to the left. However, the activity across displacement modules will be unique to a particular directional displacement in the same way that the activity across grid cell modules is unique to a location.

Now let us see how this single module network represents an example composite object such as the logo on the coffee mug. **Figure 1C** shows two points on the composite object. Each point has a location in the space of the coffee mug as well as a location in the space of the logo. In our example, suppose that attending to point *P*_1_ on the coffee mug invokes grid cell *G*_0,1_ in the coffee mug space, and attending to it on the logo invokes cell *G*_1,1_ in the logo space. The activity of these two cells will invoke displacement cell *D*_1,0_. This displacement cell will remain active, regardless of where you move on the composite object. Suppose, you switch to point *P*_2_ and this is one step below *P*_1_. Grid cells *G*_0,0_ and *G*_1,0_ will become active in the coffee mug and logo spaces, respectively. However, the relative positions of these two cells will still invoke displacement cell *D*_1,0_.

Due to the modulo operation of displacement cells it is not possible to uniquely distinguish a composite object with just one module. In our simple example, there is a 1 in 9 chance that another composite object will invoke the same displacement cell. However, with a collection of independent grid cell modules and associated displacement modules, the combined activity is highly likely to be unique. If there are *M* modules with *N* cells in each module, the chance that two composite objects will invoke an identical set of displacement cells is 1 in *N^M^*. Note that as you sense different locations on the composite object, the active grid cells will change, but the activity in the displacement cells will be stable. Thus, the overall pattern in the displacement layer is unique to a particular pair of objects, in a specific relative configuration.

A displacement layer as described above can encode relative shifts in position. In general, to completely specify a composite object, it is also necessary to encode relative scaling, rotation, and perhaps other transformations. In the coffee cup example the size and angle of the logo is just as important as the positioning. We would notice a discrepancy if the logo were half the size or tilted at 45 degrees. Residue number systems are powerful enough to handle many numerical operations. It is therefore possible that these other transformations can also be represented by operations on grid cell modules. This is an area of ongoing research for us.

**Figure 1.**
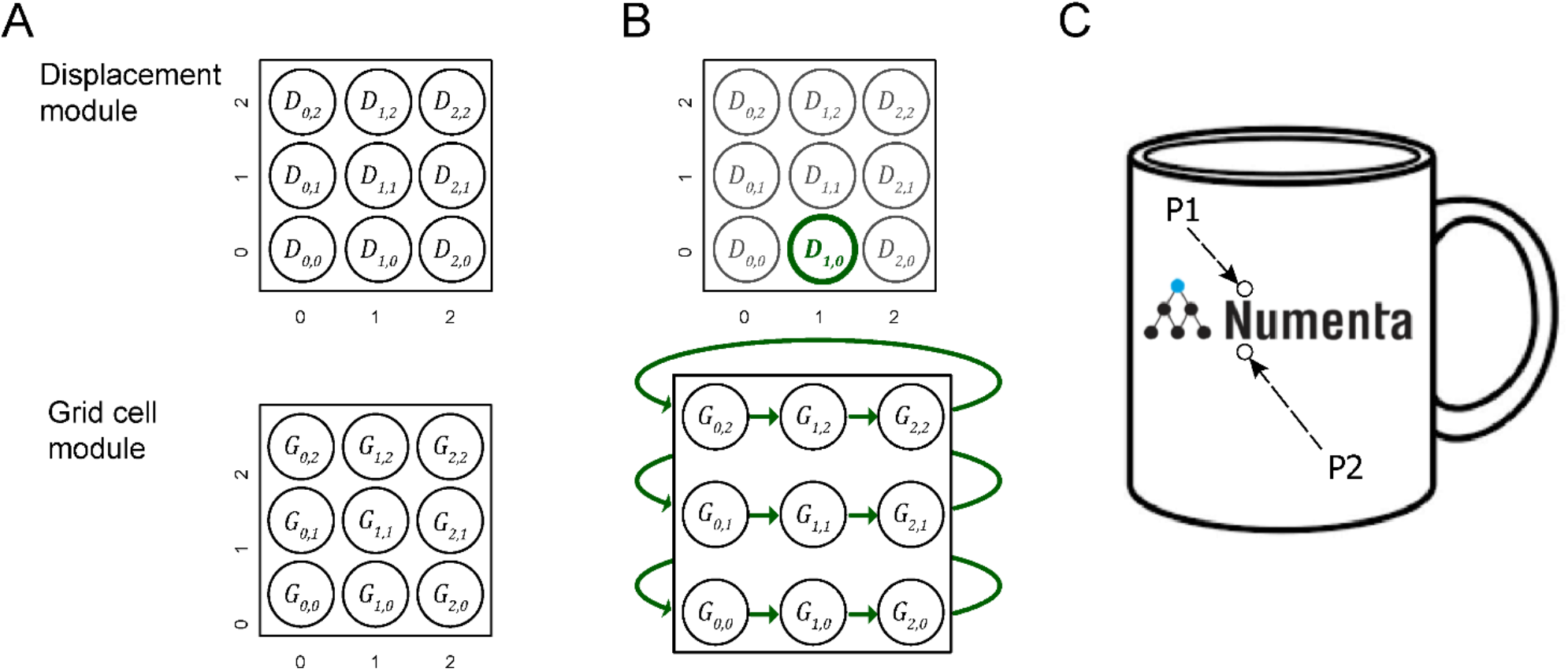
**(A)** Cells in a single grid cell module (lower square) and its corresponding displacement module (upper square). In this example each module contains 9 cells, arranged in a 3X3 lattice with associated (x,y) coordinates. Cells are arranged such that neighboring cells represent neighboring positions or displacements. For convenience we show the same number of cells in both modules. **(B)** Shows the transitions that cause displacement cell *D*_1,0_(green) to become active. Any of the 9 transitions (green arrows) will cause the cell to become active. **(C)** Two points on a coffee cup with an embedded logo. Attending to *P*_1_ in the coffee cup space followed by attending to *P*_1_ in the logo space will activate two different grid cells and a single displacement cell. If you attend to *P*_2_, in cup space and then logo space, the active grid cells will be different than with *P*_1_, but the active displacement cell will not change.

This method of representing objects allows for hierarchical composition. For example, the logo on the cup is also composed of sub-objects, such as letters and a graphic. A displacement vector placing the logo on the cup implicitly carries with it all the sub-objects of the logo. The method also allows for recursive structures. For example, the logo could contain a picture of a coffee cup with a logo. Hierarchical and recursive composition are fundamental elements of not only physical objects but language, mathematics, and other manifestations of intelligent thought. The key idea is that the identity and relative position of two previously-learned objects, even complex objects, can be represented efficiently by a single displacement vector.

### Grid Cells and Displacement Cells Perform Complementary Operations

Grid cells and displacement cells perform complementary operations. Grid cells determine a new location based on a current location and a displacement vector (i.e. movement). Displacement cells determine what displacement is required to reach a new location from a current location.

Grid cells: **(Location1 + Displacement => Location2)**

Displacement cells: **(Location2 – Location1 => Displacement)**

If the two locations are in the <u>same space</u>, then grid cells and displacement cells are useful for navigation. In this case, grid cells predict a new location based on a starting location and a given movement. Displacement cells would represent what movement is needed to get from Location1 to Location2.

If the two locations are in <u>different spaces</u> (that is the same physical location relative to two different objects) then grid cells and displacement cells are useful for representing the relative position of two objects. Grid cells convert a location in one object space to the equivalent location in a second object space based on a given displacement. In this case displacement cells represent the relative position of two objects.

We propose that grid cells and displacement cells exist in all cortical columns. They perform two fundamental and complementary operations in a location-based framework of cortical processing. By alternating between representations of locations in a single object space and representations of locations in two different object spaces, the neocortex can use grid cells and displacement cells to learn both the structure of objects and generate behaviors to manipulate those objects.

The existence of grid cells in the entorhinal cortex is well documented. We propose they also exist in all regions of the neocortex. The existence of displacement cells is a prediction introduced in this paper. We propose displacement cells are also present in all regions of the neocortex. Given their complementary role to grid cells, it is possible that displacement cells are also present in the hippocampal complex.

### Object Behaviors

Objects may exhibit behaviors. For example, consider the stapler in **Figure 4**. The top of the stapler can be lifted and rotated. This action changes the stapler’s morphology but not its identity. We don’t perceive the open and closed stapler as two different objects even though the overall shape has changed. The movement of a part of an object relative to other parts of an object is a “behavior” of the object. The behaviors of an object can be learned, and therefore they must be represented in the neural tissue of cortical columns. We can represent this behaviors in a location-based framework, again using displacement vectors. The top half and bottom half of the stapler are two components of the stapler. The relative position of the top and bottom is represented by a displacement vector in the same way as the relative position of the logo and the coffee cup. However, unlike the logo on the coffee cup, the two halves of the stapler can move relative to each other. As the stapler top rotates upward, the displacement of the stapler top to bottom changes. Thus, the rotation of the stapler top is represented by a sequence of displacement vectors. By learning this sequence, the system will have learned this behavior of the object.

**Figure 4.**
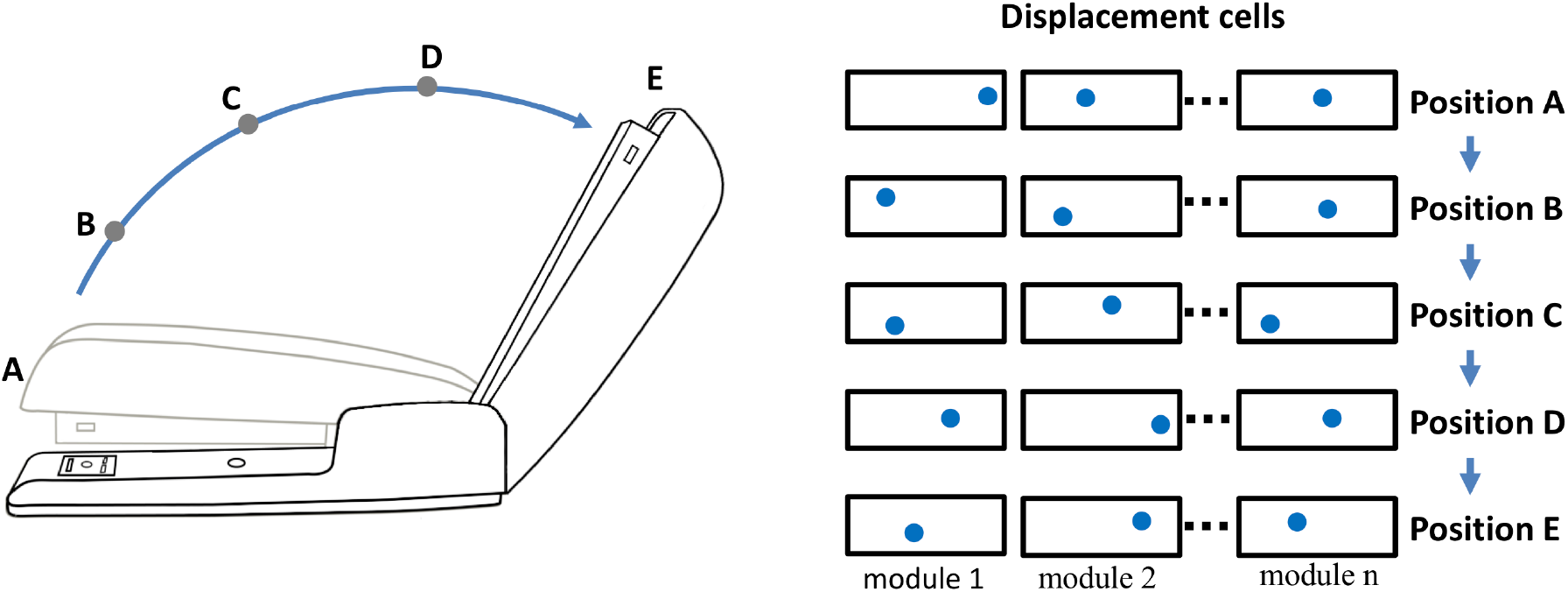
Representing Behaviors of Objects. Objects have “behaviors”, they can change their shape and features over time. The neocortex can learn these changes, but how? For example, a stapler has several behaviors, one is rotating the top relative to the base. If the top of the stapler is a component object of the stapler, with its own location space, then its position relative to the stapler base is represented by a displacement vector as illustrated in Fig. 3. (The top and base of the stapler are analogous to the logo and the cup. Unlike the logo on the cup, the location of the stapler top relative to the base can change.) The closed position is represented by displacement A and the fully open position is represented by displacement E. As the stapler top hinges from the closed to open position, the displacement vector will continually change. (Five positions, A to E, and corresponding displacement vectors are shown.) To learn this behavior, the neocortex only needs to learn the sequence of displacement vectors as the top rotates.

Opening and closing the stapler are different behaviors yet they are composed of the same displacement elements, just in reverse order. These are sometimes referred to as “high-order” sequences. Previously we described a neural mechanism for learning high-order sequences in a layer of neurons (Hawkins and Ahmad, 2016). This mechanism, if applied to the displacement modules, would allow the learning, inference, and recall of complex behavioral sequences of objects.

### “What” and “Where” Processing

Sensory processing occurs in two parallel sets of neocortical regions, often referred to as “what” and “where” pathways. In vision, damage to the “what”, or ventral, pathway is associated with the loss of ability to visually recognize objects whereas damage to the “where”, or dorsal, pathway is associated with the loss of ability to reach for an object even if it has been visually identified. Equivalent “what” and “where” pathways have been observed in other sensory modalities, thus it appears to be general principle of cortical organization (Ungerleider and Haxby, 1994; Goodale and Milner, 1992; Rauschecker, 2015). “What” and “where” cortical regions have similar anatomy and therefore we can assume they operate on similar principles.

A location-based framework for cortical function is applicable to both “what” and “where” processing. Briefly, we propose that the primary difference between “what” regions and “where” regions is that in “what” regions cortical grid cells represent locations that are allocentric, in the location space of objects, and in “where” regions cortical grid cells represent locations that are egocentric, in the location space of the body. **Figure 5** shows how a displacement vector representing movement could be generated in “what” and “where” regions. The basic operation, common to all, is that a region first attends to one location and then to a second location. The displacement cells will determine the movement vector needed to move from the first location to the second location. In a “what” region, **Figure 5C**, the two locations are in the space of an object, therefore, the displacement vector will represent the movement needed to move the finger from the first location on the object to the second location on the object. In this example, the “what” region needs to know where the finger is relative to the cup, but it does not need to know where the cup or finger is relative to the body. In a “where” region, **Figure 5B**, the two locations are in the space of the body, therefore, the displacement vector will represent how to move from one egocentric location to a second egocentric location. The “where” region can perform this calculation not knowing what object may or may not be at the second location. A more detailed discussion of processing in “where” regions is beyond the scope of this paper. We only want to point out that it is possible to understand both “what” and “where” processing using similar mechanisms by assuming different location spaces.

**Figure 5.**
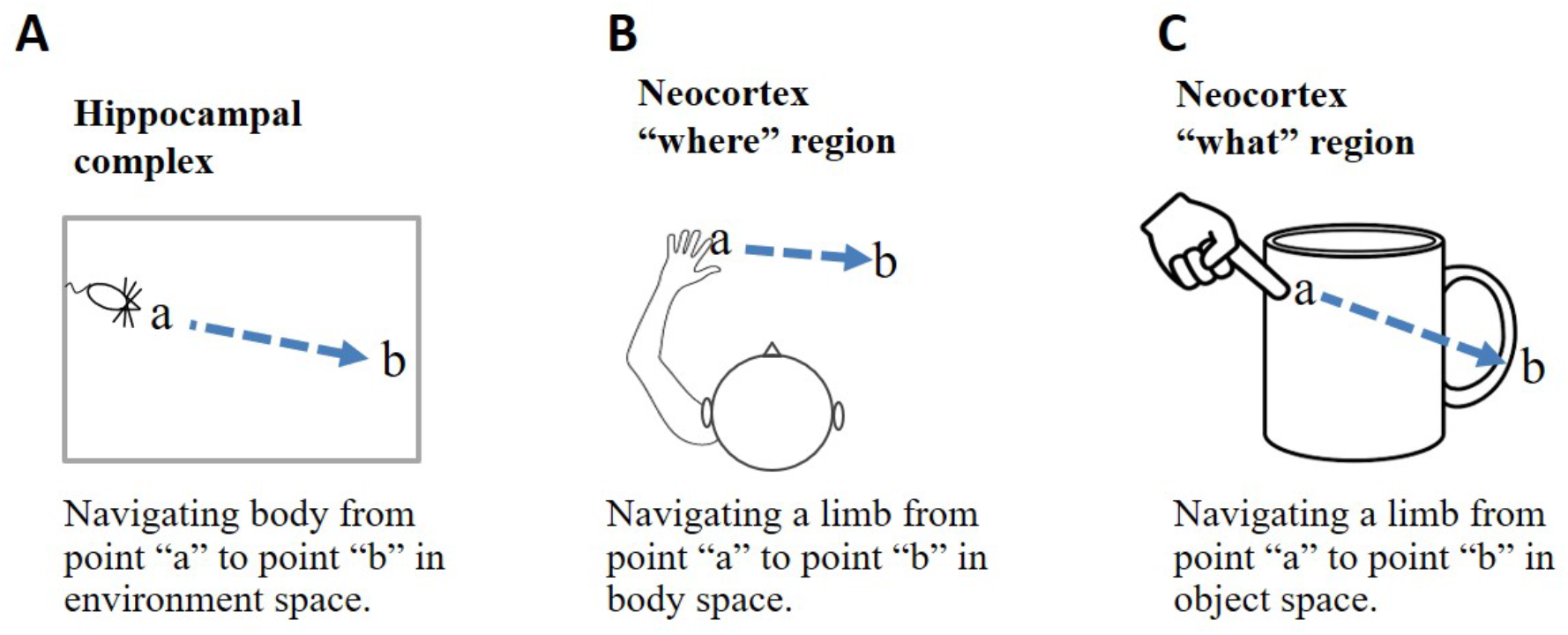
Location Processing in Different Areas of the Brain. Grid cells and displacement cells (see text) can be applied to different tasks in different areas of the brain. **(A)** If grid cell modules in the hippocampal complex are anchored by cues in an environment, then grid cell activation patterns will represent locations relative to that environment. Given two locations, **a** and **b**, displacement cells will calculate the movement vector needed to move the body from point **a** to point **b. (B)** If cortical grid cell modules are anchored relative to the body, then they will represent locations in body space. Given two locations, displacement cells will calculate the movement vector needed to move a body part from its current location to a desired new location relative to the body. **(C)** If cortical grid cell modules are anchored by cues relative to an object, then they will represent locations in the object’s space. Displacement cells will calculate the movement vector needed to move a limb or sensory organ from its current location to a new location relative to the object. Operations performed in **(B)** and **(C)** are associated with “where” and “what” regions in the neocortex.

### Rethinking Hierarchy, The Thousand Brains Theory of Intelligence

Regions of the neocortex are organized in a hierarchy (Felleman and Van Essen, 1991; Markov et al., 2014; Riesenhuber and Poggio, 1999). It is commonly believed that when sensory input enters the neocortex the first region detects simple features. The output of this region is passed to a second region that combines simple features into more complex features. This process is repeated until, several levels up in the hierarchy, cells respond to complete objects (**Figure 6A**). This view of the neocortex as a hierarchy of feature extractors also underlies many artificial neural networks (LeCun et al., 2015).

**Figure 6.**
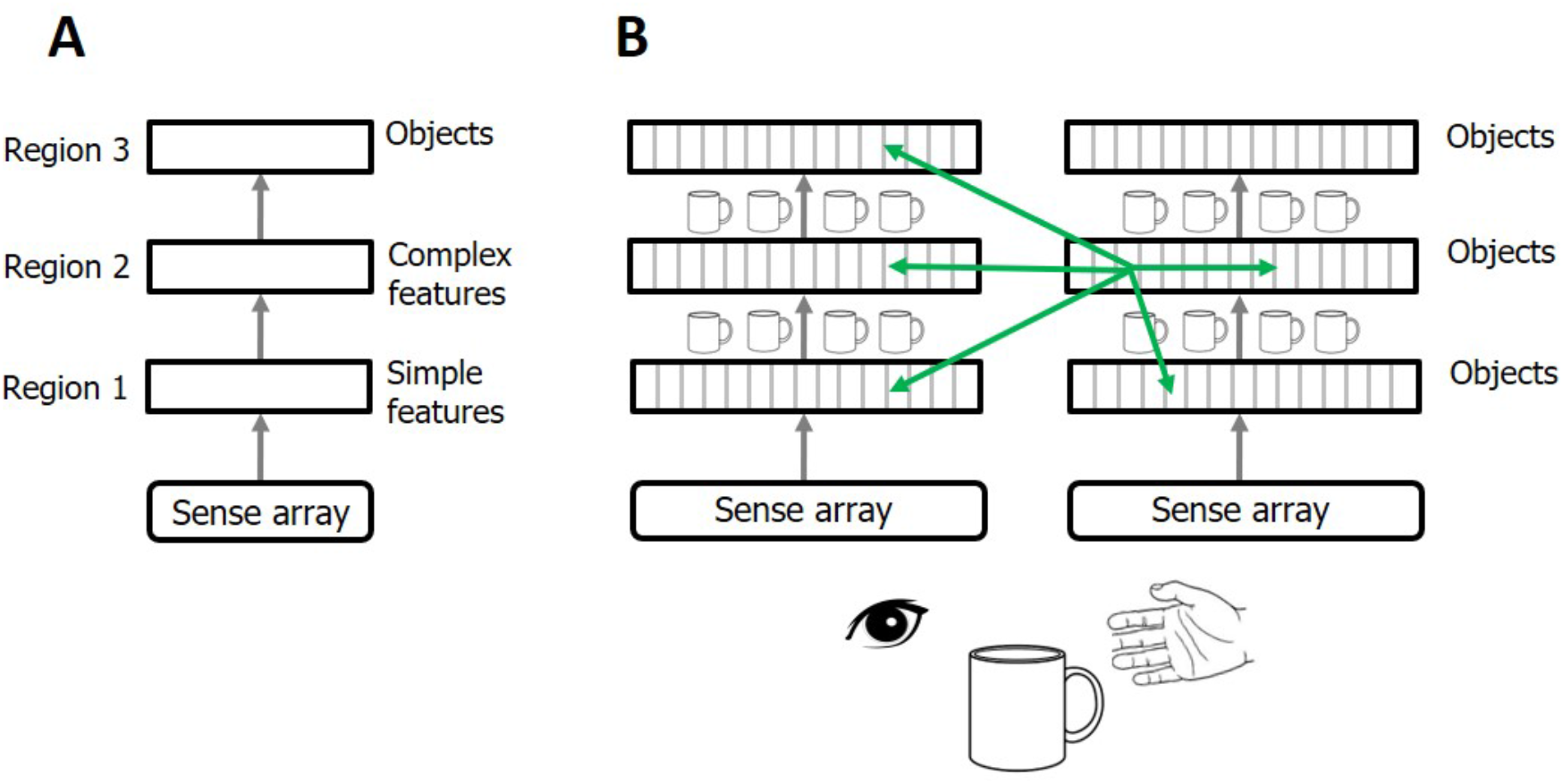
Rethinking Cortical Hierarchy **(A)** Commonly held view of cortical hierarchy. Sensory input is processed in a hierarchy of cortical regions. The first region detects simple features. The next region combines simple features into more complex features. This is repeated until a region at the top of the hierarchy forms representations of complete objects. **(B)** Modified view of cortical hierarchy. Every column in every region learns complete models of objects. (Columns learn complete models by combining sensory input with an object-centric location of that input and integrating over movements of the sensor.). Shown are two sensory hierarchies, one for vision and one for touch, both sensing the same object, a cup. There are multiple models of an object within each region, in different regions within a sensory modality, and in different sensory modalities. Although there are many models of the same object (suggested by the small cup images), the models are not identical, as each model is learned via a different subset of the sensory arrays. The green arrows denote the numerically-large cortical-cortical connections that are not hierarchical in nature. The non-hierarchical connections project within the region of origin, across hierarchical levels, across modalities, and between hemispheres. Typically, many columns will be simultaneously observing the same object. The non-hierarchical connections between columns allow them to rapidly infer the correct object (see text). Although learning objects requires movement of the sensors, inference often occurs without movement due to the non-hierarchical connections.

We propose that cortical columns are more powerful than currently believed. Every cortical column learns models of complete objects. They achieve this by combining input with a grid cell-derived location, and then integrating over movements. (See (Hawkins et al., 2017; Lewis et al., 2018) for details.) This suggests a modified interpretation of the cortical hierarchy, where complete models of objects are learned at every hierarchical level, and every region contains multiple models of objects (**Figure 6B**).

Feedforward and feedback projections between regions typically connect to multiple levels of the hierarchy. (Only one level of connection is shown in **Figure 6**.). For example, the retina projects to thalamic relay cells in LGN, which then project to cortical regions V1, V2, and V4, not just V1. This form of “level skipping” is the rule, not the exception. Therefore, V1 and V2 are both, to some extent, operating on retinal input. The connections from LGN to V2 are more divergent suggesting that V2 is learning models at a different spatial scale than V1. We predict that the spatial scale of cortical grid cells in V2 will similarly be larger than those in V1. The level of convergence of input to a region, paired with the spatial scale of its grid cells, determines the range of object sizes the region can learn. For example, imagine recognizing printed letters of the alphabet. Letters at the smallest discernable size will be recognized in V1 and only V1. The direct input to V2 will lack the feature resolution needed. However, larger printed letters would be recognized in both V1 and V2, and even larger letters may be too large for V1 but recognizable in V2. Hierarchical processing still occurs. All we are proposing is that when a region such as V1 passes information to another region such as V2, it is not passing representations of unclassified features but, if it can, it passes representations of complete objects. This would be difficult to observe empirically if objects are represented by population codes as proposed in (Hawkins et al., 2017). Individual neurons would participate in many different object representations and if observed in isolation will appear to represent sensory features, not objects. The number of objects that a cortical column can learn is large but limited (Hawkins et al., 2017). Not every column can learn every object. Analysis of system capacity requires a more thorough understanding of hierarchical flow and is beyond the scope of this paper.

There are many cortical-cortical projections that are inconsistent with pure hierarchical processing (**Figure 6B green arrows**). For example, there are long range projections between regions in the left and right hemispheres (Clarke and Zaidel, 1994), and there are numerous connections between regions in different sensory modalities, even at the lowest levels of the hierarchy (Suter and Shepherd, 2015; Driver and Noesselt, 2008; Schroeder and Foxe, 2005). These connections may not be hierarchical as their axons terminate on cells located outside of cellular layers associated with feedforward or feedback input. It has been estimated that 40% of all possible region-to-region connections actually exist which is much larger than a pure hierarchy would suggest (Felleman and Van Essen, 1991). What is the purpose of these long-range non-hierarchical connections? In (Hawkins et al., 2017) we proposed that cell activity in some layers (e.g. L4 and L6) of a column changes with each new sensation, whereas, cell activity in other layers (e.g. L2/3), representing the observed “object”, are stable over changing input. We showed how long-range associative connections in the “object” layer allow multiple columns to vote on what object they are currently observing. For example, if we see and touch a coffee cup there will be many columns simultaneously observing different parts of the cup. These columns will be in multiple levels of both the visual and somatosensory hierarchies. Every one of these columns has a unique sensory input and a unique location, and therefore, long-range connections between the cells representing location and input do not make sense. However, if the columns are observing the same object, then connections between cells in the object layer allow the columns to rapidly settle on the correct object. Thus, non-hierarchical connections between any two regions, even primary and secondary sensory regions in different sensory modalities, make sense if the two regions often observe the same object at the same time. See (Hawkins et al., 2017) for details.

One of the classic questions about perception is how does the neocortex fuse different sensory inputs into a unified model of a perceived object. We propose that the neocortex implements a decentralized model of sensor fusion. For example, there is no single model of a coffee cup that includes what a cup feels like and looks like. Instead there are hundreds of models of a cup. Each model is based on a unique subset of sensory input within different sensory modalities. There will be multiple models based on visual input and multiple models based on somatosensory input. Each model can infer the cup on its own by observing input over movements of its associated sensors. However, long-range non-hierarchical connections allow the models to rapidly reach a consensus of the identity of the underlying object, often in a single sensation.

Just because each region learns complete models of objects does not preclude hierarchical flow. The main idea is that the neocortex has hundreds, likely thousands, of models of each object in the world. The integration of observed features does not just occur at the top of the hierarchy, it occurs in every column at all levels of the hierarchy. We call this “The Thousand Brains Theory of Intelligence.”

## Discussion

In 1979 Francis Crick wrote an essay titled, “Thinking about the Brain” (Crick, 1979). In it he wrote, “In spite of the steady accumulation of detailed knowledge, how the human brain works is still profoundly mysterious.” He posited that over the coming years we would undoubtedly accumulate much more data about the brain, but it may not matter, as “our entire way of thinking about such problems may be incorrect.” He concluded that we lacked a “theoretical framework”, a framework in which we can interpret experimental findings and to which detailed theories can be applied. Nearly forty years after Crick wrote his essay, his observations are still largely valid.

Arguably, the most progress we have made towards establishing a theoretical framework is based on the discovery of place cells and grid cells in the hippocampal complex. These discoveries have suggested a framework for how animals learn maps of environments, and how they navigate through the world using these maps. The success of this framework has led to an explosion of interest in studying the entorhinal cortex and hippocampus.

In this paper we are proposing a theoretical framework for understanding the neocortex. Our proposed cortical framework is a derivative of the framework established by grid cells and place cells. Mechanisms that evolved for learning the structure of environments are now applied to learning the structure of objects. Mechanisms that evolved for tracking the location of an animal in its environments are now applied to tracking the location of limbs and sensory organs relative to objects in the world. How far this analogy can be taken is uncertain. Within the circuits formed by the hippocampus, subiculum, and entorhinal cortex are grid cells (Hafting et al., 2005), place cells (O’Keefe and Dostrovsky, 1971; O’Keefe and Burgess, 2005), head direction cells (Taube et al., 1990; Giocomo et al., 2014; Winter et al., 2015), border cells (Lever et al., 2009), object vector cells (Deshmukh and Knierim, 2013) and others, plus many conjunctive cells that exhibit properties that are combinations of these (Sargolini et al., 2006; Brandon et al., 2011; Stensola et al., 2012; Hardcastle et al., 2017). We are currently exploring the idea that the neocortex contains cells that perform equivalent functions to the variety of cells found in the hippocampal complex. The properties of these cells would only be detectable in an awake animal actively sensing learned objects. The recent work of (Long and Zhang, 2018) suggests this might be true.

### Orientation

In the entorhinal cortex, and elsewhere in the brain, are found head direction cells (Sargolini et al., 2006; Raudies et al., 2016; Taube et al., 1990). These cells represent the allocentric orientation of an animal relative to its environment. Inferring where you are via sensation, predicting what you will sense after moving, and determining how to move to get to a new location all require knowing your current orientation relative to your environment. In the models reviewed in (Hasselmo, 2009; Hasselmo et al., 2010) head direction cells are critical for accurately transitioning between spatial locations. The same need for orientation exists throughout the neocortex. For example, knowing that a finger is at a particular location on a coffee cup is not sufficient. The finger also has an orientation relative to the cup (which way it is rotated and its angle at contact). Predicting what the finger will sense when it contacts the cup or what movement is required to reach a new location on the cup requires knowing the finger’s orientation relative to the cup in addition to its location. Therefore, we predict that within each cortical column there will be a representation of orientation that performs an analogous function to head direction cells in the hippocampal complex. How orientation is represented in the cortex is unknown. There could be a set of orientation cells each with a preferred orientation, similar to head direction cells, but we are not aware of any evidence for this. Alternately, orientation could be represented via a population code, which would be more difficult to detect. For example, in somatosensory regions orientation could be represented by activating a sparse subset of egocentric orientation detectors (Hsiao et al., 2002; Bensmaia et al., 2008; Pruszynski and Johansson, 2014). How orientation is represented and interacts with cortical grid cells and displacement cells is largely unknown. It is an area we are actively studying.

### Prediction

A long standing principle behind many theories of cortical function is prediction (Lashley, 1951; Hawkins and Blakeslee, 2004; Lotter et al., 2018). By representing the location of a sensor, a cortical column can associate sensory information within the location space of each object, similar to the way place cells associate sensory information with locations (Komorowski et al., 2009). This enables a column to build powerful predictive models. For example, when moving your finger from the bottom of a cup to the top, it can predict the sensation regardless of how the cup is rotated with respect to the sensor. Representing composite objects using displacement cells enables a column to generalize and predict sensations even when encountering a novel object. For example, suppose we see a cup with a familiar logo (**Figure 3A**) and that portions of the logo are obscured. Once a column has recognized the logo and the cup, it can make predictions regarding the entire logo in relation to the cup even if that combined object is new. Building such predictive models would be much harder without an explicit representation of location. In previous papers we proposed dendritic mechanisms that could serve as the neural basis for predictive networks (Hawkins and Ahmad, 2016; Hawkins et al., 2017). Overall, prediction underlies much of the framework discussed in this paper.

### Attention

One of the key elements of a location-based framework for cortical processing is the ability of an area of cortex to rapidly switch between object spaces. To learn there is a logo on the coffee cup we need to alternate our attention between the cup and the logo. With each shift of attention, the cortical grid cells re-anchor to the location space of the newly attended object. This shift to a new object space is necessary to represent the displacement between two objects, such as the logo and the cup. It is normal to continuously shift our attention between the objects around us. With each newly attended object the cortical grid cells re-anchor in the space of the new object, and displacement cells represent where the new object is relative to the previously attended object. Changing attention is intimately tied to movement of the sensor, re-anchoring of grid cells, and, as widely believed, feedback signals to the thalamus (Crick, 1984; McAlonan et al., 2006), presumably to select a subset of input for processing. How these elements work together is poorly understood and represents an area for further study.

### Uniqueness of Location Code

Our proposal is based on the idea that a set of grid cell modules can encode a very large number of unique locations. There are some observations that suggest that grid cells, on their own, may not be capable of forming enough unique codes. For example, because each grid cell exhibits activity over a fairly large area of physical space (Hafting et al., 2005), the activation of the cells in a grid cell module is not very sparse. Sparsity is helpful for creating easily discernable unique codes. The lack of sparsity can be overcome by sampling the activity over more grid cell modules, but not enough is known about the size of grid cell modules and how many can be realistically sampled. (Gu et al., 2018) have shown that grid cell modules are composed of smaller sub-units that activate independently, which would also increase the representation capacity of grid cells. Another factor impacting capacity is conjunctive cells. In the entorhinal cortex there are more conjunctive cells than pure grid cells. Conjunctive cells exhibit some combination of “gridness” plus orientation and/or other factors (Sargolini et al., 2006). Conjunctive cells may have a sparser activation than pure grid cells and therefore would be a better basis for forming a set of unique location codes. If the neocortex has cells similar to conjunctive cells, they also might play a role in location coding. Not enough is known about how grid cells, orientation cells, and conjunctive cells work together to suggest exactly how locations are encoded in the neocortex. As we learn more about location coding in the neocortex, it is important to keep these possibilities in mind.

### Where are Grid Cells and Displacement Cells in the Neocortex?

The neocortex is commonly divided into 6 layers that run parallel to the surface. There are dozens of different cell types, therefore, each layer contains multiple cell types. Several lines of evidence suggest that cortical grid cells are located in L6 (specifically L6 cortical-cortical neurons (Thomson, 2010)) and displacement cells are located in L5 (specifically L5 thick-tufted neurons) (**Fig. 7**).

**Figure 7.**
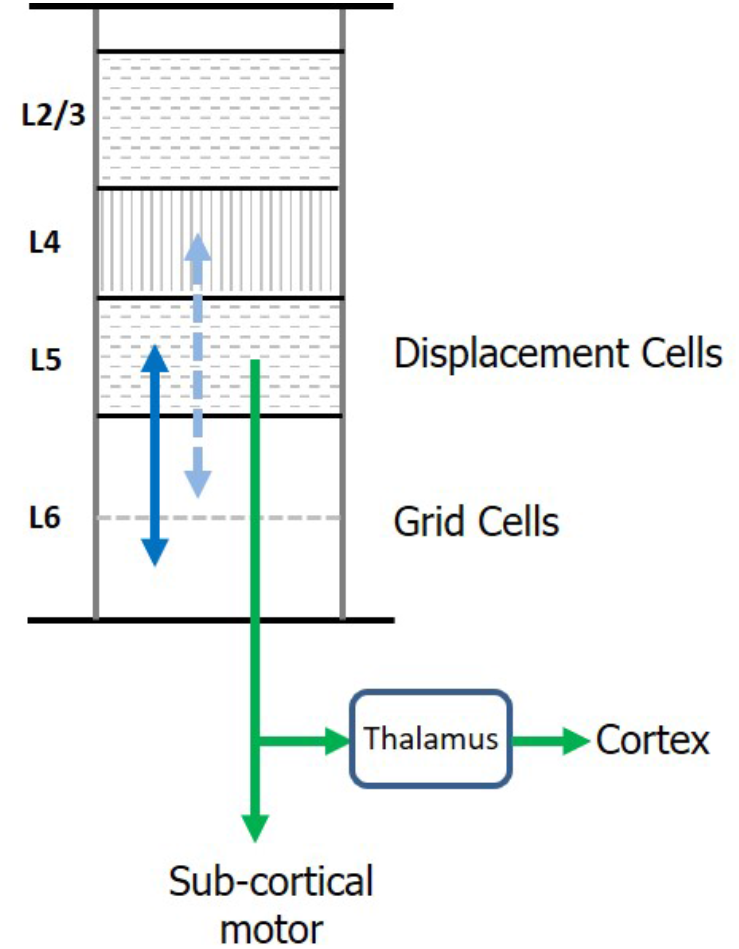
Location of Grid Cells and Displacement Cells in the Neocortex. The neocortex contains dozens of cell types commonly organized into six cellular layers. Here we show a simple drawing of a cortical column. We propose cortical grid cells are located in layer 6 and displacement cells are in layer 5. A requirement of our proposal is that cortical grid cells make bi-lateral connections with displacement cells (solid blue line). Another requirement is that, when combined with a representation of orientation, they make bilateral connections with cells in layer 4 (dashed blue line). This is how the column predicts the next input into layer 4. Displacement cells match the unusual connectivity of layer 5 “thick tufted” neurons, which are the motor output cells of the neocortex. These neurons send their axon down into the white matter where the axon splits (green arrows). One branch terminates in sub-cortical structures responsible for motor behavior. The second axon branch terminates on relay cells in the thalamus which become the feedforward input to a hierarchically-higher cortical region. As explained in the text, displacement cells can alternate between representing movements and representing the composition of multiple objects. We propose that L5 thick tufted cells alternate between these two functions which aligns with their unusual connectivity.

One piece of evidence suggesting cortical grid cells are in L6 is the unusual connectivity between L4 and L6. L4 is the primary input layer. However, feed forward input forms less than 10% of the synapses on L4 cells (Ahmed et al., 1994, 1997; Sherman and Guillery, 2013), whereas approximately 45% of the synapses on L4 cells come from L6a cortical-cortical neurons (Ahmed et al., 1994; Binzegger et al., 2004). Similarly, L4 cells make large numbers of synapses onto those same L6 cells (McGuire et al., 1984; Binzegger et al., 2004; Kim et al., 2014). Also, the connections between L6 and L4 are relatively narrow in spread (Binzegger et al., 2004). The narrow connectivity between L6 and L4 is reminiscent of the topologically-aligned bidirectional connectivity between grid cells in MEC and place cells in hippocampus (Zhang et al., 2013; Rowland et al., 2013). We previously showed how the reciprocal connections between L6 and L4 can learn the structure of objects by movement of sensors if L6 represents a location in the space of the object (Lewis et al., 2018). For a column to learn the structure of objects in this fashion requires bidirectional connections between cells receiving sensory input and cells representing location. L6a is the only known set of cells that meet this requirement. Also, grid cells use motor input to update their representations for path integration. Experiments show significant motor projections to L6 (Nelson et al., 2013; Leinweber et al., 2017). The current experimental evidence for the presence of grid cells in the neocortex is unfortunately mute on what cortical layers contain grid cells. It should be possible to experimentally determine this in the near future. Our prediction is they will be in L6.

The main evidence for displacement cells being in L5 is again connectivity. A subset of L5 cells (known as “L5 thick-tufted cells”) that, as far as we know, exists in all cortical regions, projects sub-cortically to brain regions involved with motor behavior. (For example, L5 cells in the visual cortex project to the superior colliculus which controls eye movements.) These L5 cells are the motor output cells of the neocortex. However, the same L5 cells send a branch of their axon to thalamic relay nuclei, which then project to hierarchically higher cortical regions (Douglas and Martin, 2004; Guillery and Sherman, 2011; Sherman and Guillery, 2011). It is difficult to understand how the same L5 cells can be both the motor output and the feedforward input to other regions. One interpretation put forth by Guillery and Sherman is that L5 cells represent a motor command and that the feedforward L5 projection can be interpreted as an efference copy of the motor command (Guillery and Sherman, 2011, 2002).

We offer a possible alternate interpretation. The L5 cells in question are displacement cells and they alternately represent movements (sent sub-cortically) and then represent compositional objects (sent to higher regions via thalamic relay cells). As described above, displacement cells will represent a movement vector when comparing two locations in the same space and will represent composite objects when comparing two locations in two different spaces. These two rapidly-changing representations could be disambiguated at their destination either by phase of an oscillatory cycle or by physiological firing patterns (Hasselmo and Brandon, 2012; Hasselmo, 2008; Burgess et al., 2007). Although we are far from having a complete understanding of what the different cellular layers do and how they work together, a location-based framework offers the opportunity of looking anew at the vast body of literature on cortical anatomy and physiology and making progress on this problem.

### Location-based Framework for High-level Thought and Intelligence

We have described our location-based framework using examples from sensory inference. Given that the anatomy in all cortical regions is remarkably similar, it is highly likely that everything the neocortex does, including language and other forms of high-level thought, will be built upon the same location-based framework. In support of this idea, the current empirical evidence that grid cells exist in the neocortex was collected from humans performing what might be called “cognitive tasks”, and it was detected in cortical regions that are far from direct sensory input. (Doeller et al., 2010; Constantinescu et al., 2016; Jacobs et al., 2013).

The location-based framework can be applied to physical structures, such as a cup, and to abstract concepts, such as mathematics and language. A cortical column is fundamentally a system for learning predictive models. The models are learned from inputs and movements that lead to changes in the input. Successful models are ones that can predict the next input given the current state and an anticipated movement. However, the “inputs” and “movements” of a cortical column do not have to correspond to physical entities. The “input” to a column can originate from the retina or it can originate from other regions of the neocortex that have already recognized a visual object such as a word or a mathematical expression. A “movement” can represent the movement of the eyes or it can represent an abstract movement, such as a verb or a mathematical operator.

Success in learning a predictive model requires discovering the correct dimensionality of the space of the object, learning how movements update locations in that space, and associating input features with specific locations in the space of the object. These attributes apply to both sensory perception and high-level thought. Imagine a column trying to learn a model of a cup using visual input from the retina and movement input from a finger. This would fail, as the location spaced traversed by the finger would not map onto the feature space of the object as evidenced by the changing inputs from the eyes. Similarly, when trying to understand a mathematical problem you might fail when using one operator to manipulate an equation but succeed by switching to a different operator.

Grid cells in the neocortex suggests that all knowledge is learned and stored in the context of locations and location spaces and that “thinking” is movement through those location spaces. We have a long way to go before we understand the details of how the neocortex performs cognitive functions, however, we believe that the location-based framework will not only be at the core of the solutions to these problems, but will suggest solutions.

## Conclusion

It is sometimes said that neuroscience is “data rich and theory poor”. This notion is especially true for the neocortex. We are not lacking empirical data as much as lacking a theoretical framework that can bridge the gap between the heterogeneous capabilities of perception, cognition, and intelligence and the homogeneous circuitry observed in the neocortex. The closest we have to such a framework today is hierarchical feature extraction, which is widely recognized as insufficient.

One approach to developing a theory of neocortical function is to build in-silico models of a cortical column based on detailed anatomical data (Helmstaedter et al., 2007; Markram et al., 2015). This approach starts with anatomy and hopes to discover theoretical principles via simulation of a cortical column. We have used a different method. We start with a detailed function that we know the neocortex performs (such as sensory-motor learning and inference), we deduce neural mechanisms that are needed to perform those functions (such as cells that represent location), and then map those neural mechanisms onto detailed biological data.

Based on this method, this paper proposes a new framework for understanding how the neocortex works. We propose that grid cells are present everywhere in the neocortex. Cortical grid cells track the location of inputs to the neocortex in the reference frames of the objects being observed. We propose the existence of a new type of neuron, displacement cells, that complement grid cells, and are similarly present throughout the neocortex. The framework shows how it is possible that a small patch of cortex can represent and learn the morphology of objects, how objects are composed of other objects, and the behaviors of objects. The framework also leads to a new interpretation of how the neocortex works overall. Instead of processing input in a series of feature extraction steps leading to object recognition at the top of the hierarchy, the neocortex consists of thousands of models operating in parallel as well as hierarchically.

Introspection can sometimes reveal basic truths that are missed by more objective experimental techniques. As we go about our day we perceive thousands of objects, such as trees, printed and spoken words, buildings, and people. Everything is perceived at a location. As we attend to each object we perceive the distance and direction from ourselves to these objects, and we perceive where they are relative to each other. The sense of location and distance is inherent to perception, it occurs without effort or delay. It is self-evident that the brain must have neural representations for the locations of objects and for the distances between the objects as we attend to them in succession. The novelty of our claim is that these locations and distances are calculated everywhere in the neocortex, they are the principal data types of cortical function, perception, and intelligence.

## Acknowledgements

We thank David Eagleman, Weinan Sun, Michael Hasselmo, and Mehmet Fatih Yanik for their feedback and help with this manuscript. We also thank numerous collaborators at Numenta over the years for many discussions, especially Donna Dubinsky, Christy Maver, Celeste Baranski, Luiz Scheinkman, and Teri Fry.

